# Improved consistency in estimates of conditional false discovery rates increases power relative to both existing methods and parametric estimators

**DOI:** 10.1101/414326

**Authors:** James Liley, Chris Wallace

## Abstract

A common aim in high-dimensional association studies is the identification of the subset of investigated variables associated with a trait of interest. Using association statistics on the same variables for a second related trait can improve power. An important quantity in such analyses is the conditional false-discovery rate (cFDR), the probability of non-association with the trait of interest given p-value thresholds for both traits. The cFDR can be used for hypothesis testing and as a posterior probability in its own right. In this paper, we propose new estimators for the cFDR based on kernel density estimates and mixture-Gaussian models of effect sizes, the latter also allowing estimation of a ‘local’ form of cFDR (cfdr). We also propose a general non-parametric improvement to existing estimators based on estimating a posterior probability previously estimated at 1. We find that new estimators have the desirable property of smooth rejection regions, but, unexpectedly, do not improve the power of the method, even when distributional assumptions are true. Furthermore, we find that although the local cfdr represents a theoretically optimal decision boundary, noisiness in its estimation means it is less powerful than corresponding cFDR estimates. We find, however, that the non-parametric adjustment increases power for every estimator. We demonstrate the best method on transcriptome-wide association study datasets for breast and ovarian cancers. The findings from this analysis are of both theoretical and pragmatic interest, giving insight into the nature of cFDR and the behaviour of false-discovery rates in a two-dimensional setting. Our methods allow improved control over the behaviour of the cFDR estimator and improved power in high-dimensional hypothesis testing.

## 1 Introduction

In the past decade, the progress of biological investigation has been characterised by increasing size of datasets and increasing diversity and precision of phenotypes under investigation. This suggests ‘leverage’ approaches, in which we seek to learn more about one trait by using data from another. Specifically, given a set of p-values *P* from association tests for a set of hypotheses for a trait under investigation, and corresponding values *Q* of some covariate which has different distributions amongst associations and non-associations with *P*, we seek to use the values of *Q* to strengthen our ability to find associations with *P*.

A range of methods for this problem have been proposed. In a Bayesian setting, Ferkingstad *and others* (2008) determine posterior probabilities of association for each hypothesis using a frequentist p-value for the *i*th hypothesis and a prior probability of association modulated according to the covariate, with values ‘binned’ into categories. A frequentist approach by Ignatiadis *and others* (2016) weights each hypothesis under consideration using the covariate value for that hypothesis, with a form of the Benjamini-Hochberg procedure used on resultant weighted observations. A third approach by Zablocki *and others* (2014) parametrises the joint distribution of the covariate and p-values allowing continuous modulation of the probability of association with changes in the covariate. This enables estimation of the probability of association for a hypothesis of interest given p-values for both the trait of interest and the covariate, a quantity which we will revisit in this work.

The method we will primarily consider in this paper is termed the conditional false-discovery rate (cFDR). First proposed in Andreassen *and others* (2013), the cFDR represents the probability of non-association with a phenotype of interest (we will denote this event 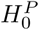) given p-value thresholds *p, q* on observed p-values *P, Q* for both the trait under investigation and a second trait (*Q*):

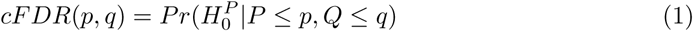

When estimated for each hypothesis at the observed values of *p, q*, the cFDR can be used for hypothesis testing. The cFDR has been used successfully to lever secondary datasets in genome-wide association studies (GWAS) (Andreassen *and others*, 2014; Andreasson *and others*, 2014; Liley and Wallace, 2015). The cFDR is powerful in many cases and effective in its simplicity and interpretability.

None of the above methods is clearly dominant in every case. In general, the problem is complex: the optimal method depends on the distribution of effect sizes for the trait under investigation, the type and distribution of covariate, and the number of variables. Furthermore, the best method depends on the desired outcome and measure of type-1 error rate.

In a recent work (Liley and Wallace, 2018), we established a method to control FDR when using cFDR for association testing. While FDR control is assured, the existing method has the shortcoming that rejection regions are often highly discontinuous and dependent on ordering of the p-values. Additionally, estimation of the ‘cfdr’ statistic used by Zablocki *and others* (2014):

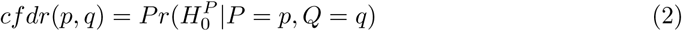

is not supported. Because our FDR controlling method is immediately adaptable to different estimators of cFDR, there is scope to diversify the range of cFDR associated methods to overcome these shortcomings and adapt to different problems.

In this work, we examine new estimators for cFDR, and evaluate their performance on simulated data. We also find that although the cfdr can be used to construct optimal rejection regions, its estimation is so noisy that it may often be less effective than the cFDR in practice. Finally, we introduce a non-parametric improvement to the existing cFDR method, and show this leads to a substantial improvement in power over existing methods. We apply the best-performing method to datasets from transcriptome-wide association studies (TWAS) as an example.

## 2 Overview of existing estimator and FDR control

Suppose we have a set of p-value pairs (*p_i_, q_i_*) where *p*_1_, *p*_2_, …, *p_n_* are p-values from association tests in a study of interest considered to be draws from the random variable *P*, and *q*_1_, *q*_2_, …, *q_n_* are corresponding p-values from the second study, considered to be draws from *Q*.

The current estimator cFDR uses a multiset of p-value pairs *X*, which may or may not be the set *S*. We expand definition (1) as

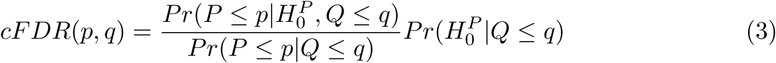

Under the assumption 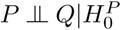, we have 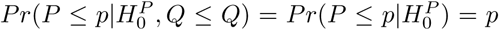. The two remaining quantities (which we will focus on in this paper) are approximated as

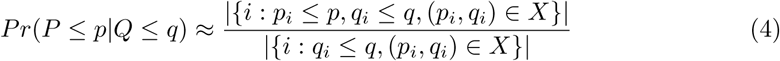

and

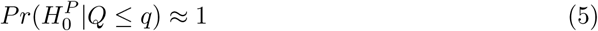

giving the estimate

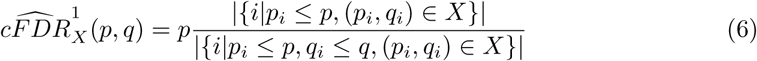

noting the dependence on *X*, and noting by the superscript ‘1’ the original estimator. Estimate (4) is generally consistent, and estimate (5) is conservative, meaning that 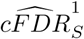 is asymptotically conservative in that under reasonably general circumstances

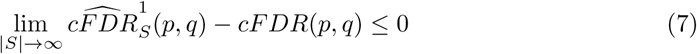

for (*p, q*) ∈ (0, 1)^2^.

The FDR-controlling method we use here begins by dividing hypotheses into ‘folds’. We then construct ‘L-regions’ for each observation (*p_i_, q_i_*) ∈ *S* which roughly represent the set of points (*p, q*) for which 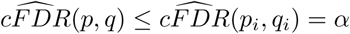. Strictly, we define

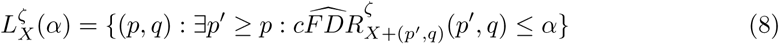

where *ζ* indicates the type of cFDR estimator used (so far we have only introduced one estimator in (6) but others will be proposed in Section 3). The L-region for point *p_i_, q_i_* is computed leaving out the points in the same fold as *p_i_, q_i_*; that is, using *X* = *S* − (fold containing *i*).

The PDF *f*_0_ of 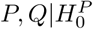 is then estimated, using the fact that 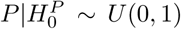 and 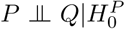, assuming 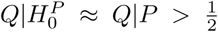 and assuming a mixture-Gaussian model for 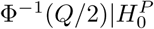 (this assumption is discussed in a later section of this paper). For each such region L, we then define ‘v-values’ (an analogue of p-values) as

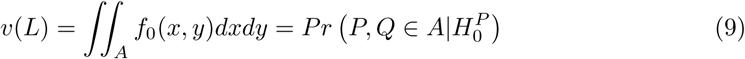

and the Benjamini-Hochberg (Benjamini and Hochberg, 1995) method is applied to control FDR. Because the FDR-controlling method uses only regions *L* rather than actual values of 7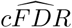, the values 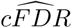 can be scaled arbitrarily and the results will not change. Importantly, FDR control will be achieved even if the estimate 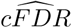 is non-conservative. For technical details regarding FDR control, we refer the reader to Liley and Wallace (2018).

When used either as a hypothesis test or as a direct estimate, 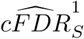 can be a poor approximation of the quantity *Pr*(*P* ≤ *p*|*Q* ≤ *q*) in regions near the *P* = 0 and *Q* = 0 boundaries of the unit square. At these extremes, the approximation in (4) has large discontinuities around observed *p_i_, q_i_*. This is demonstrated in figure 1, which shows that 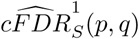 can vary twofold in any neighbourhood of certain *p_i_, q_i_* ∈ *S*. Such extreme variation at a small scale is undesirable as very small fluctuations in observed data should not have marked effects on test statistics. It is important to note that a similar problem affects posterior estimates of FDR in the Bayesian sense (Efron *and others*, 2008), and in the Benjamini-Hochberg method itself, in which the acceptance or rejection of any null is dependent on where it falls in the order of other P-values. We propose methods involving parametrisation or smoothing of the bivariate distribution of (*P, Q*), averting the problem shown in figure 1. We also introduce an approximation of the quantity 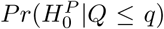 in equation (5), making the estimator of cFDR closer to consistent (up to the factor of 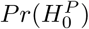, and assuming the consistency of the estimate of 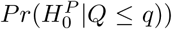 rather than conservative. We show this improves the power of the method.

**Figure 1:**
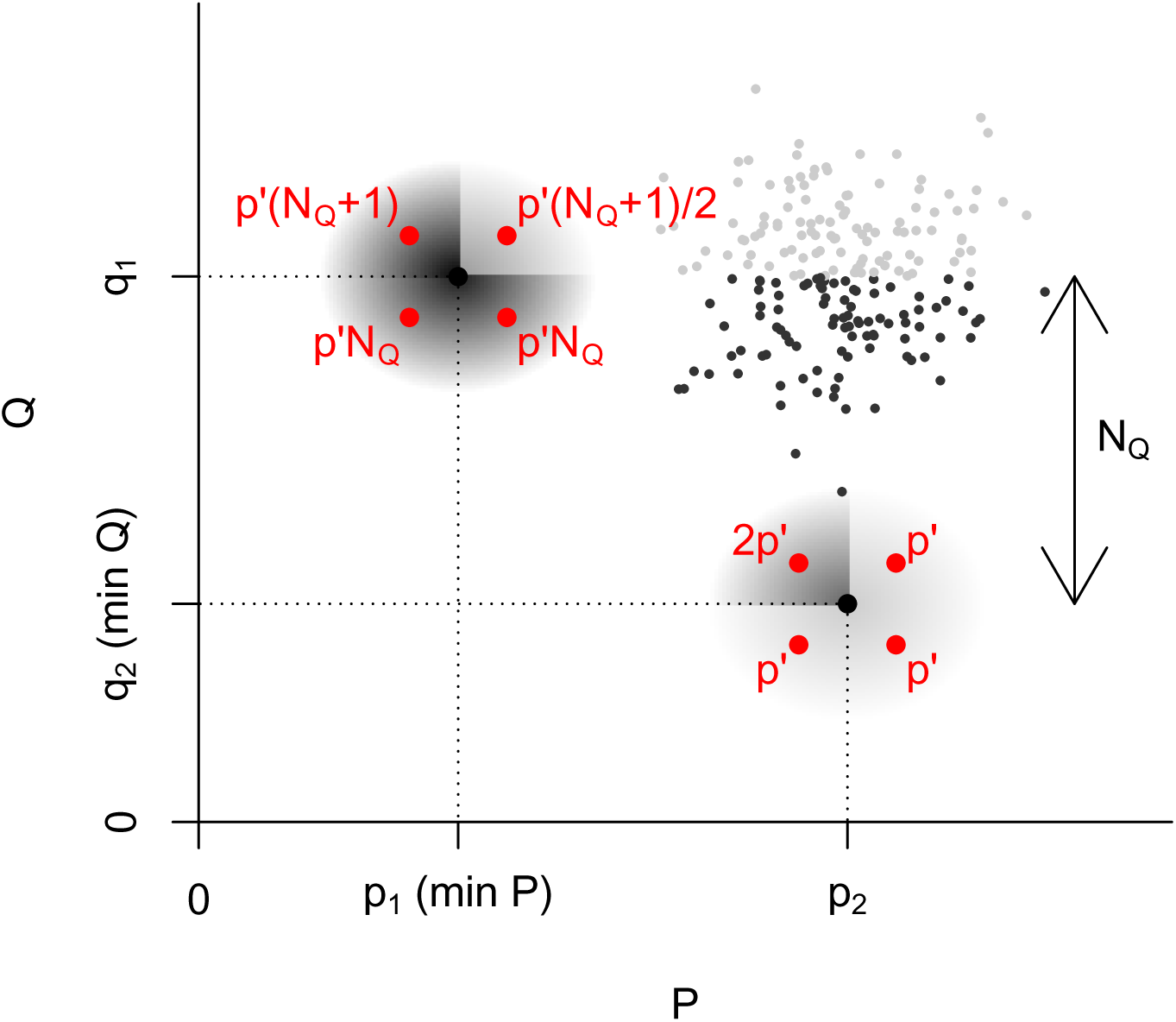
Dependence of 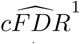 values on location of nearby points. In this example, we denote by (*p*_1_, *q*_1_), (*p*_2_, *q*_2_) the points at the (unique) left and lower extremes of the observed p-value distribution respectively; that is, *p*_1_ = min(*p_i_*), *q*_2_ = min(*q_i_*). We set *N_Q_* as the number of points with *q*_1_≤ *q_i_* ≤ *q*_2_ (small black points). If we add a test point (*p′, q′*) (shown in red) in a small neighbourhood of either (*p*_1_, *q*_1_) or (*p*_2_, *q*_2_), the estimated 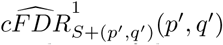 (shown in red next to the point) differs by a factor of 2 in different quadrants of the neighbourhood.

## 3 New metrics for association

### 3.1 Alternative estimators for cFDR

We now move to consider alternative estimators for cFDR. The estimate 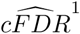 is based on empirical quantities estimated directly by counting points (ie, empirical estimation of the joint CDF of (*P, Q*)) as per definition (6), noting the dependence on the set *S*. We consider here estimators based on approximating the joint distribution of *P, Q*: one usinga kernel density estimator (KDE) and one using a bivariate mixture-normal parametrisation. These estimators enforce continuity of 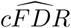 on the open unit square, and thus are robust to small deviations in p-values, overcoming the effect detailed in figure 1. It is easiest to visualise parametrisations as distributions over the unsigned Z scores (*Z_p_, Z_q_*) = (−Φ^−1^(*P*/2), −Φ^−1^(*Q*/2)) with Φ^−1^ (*x*) denoting the standard normal quantile function at *x*.

To avoid distributional assumptions while maintaining a smooth form for the density of *P, Q*, a second estimator of *Pr*(*P* < *p*|*Q* < *q*) can be derived from a two-dimensional kernel density (KDE). We had no reason to prefer any kernel function over another, so opted to use a normal kernel with constant variance *I*_2_. The PDF corresponding to *Z_p_, Z_q_* at *x, y* was modelled in the usual way as

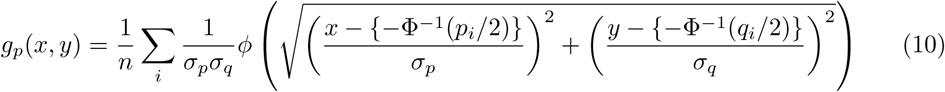

where *ϕ* (.) is the standard normal density. Values *σ_p_* and *σ_q_* are determined using a standard method based on the observations *p_i_, q_i_* ∈ *X* (Sheather and Jones, 1991).

We denote the estimator of cFDR derived from the KDE for an arbitrary point (*p, q*) with Z scores (*z_p_, z_q_*) = (Φ^−1^(*p*/2), Φ^−1^(*q*/2)) as

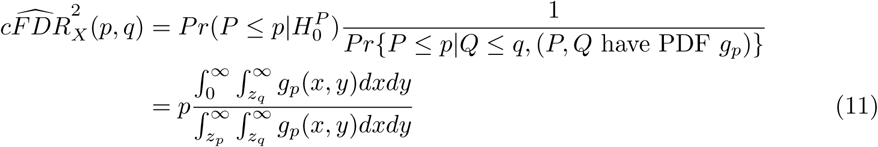

where *X* in this case refers to the set of points used in fitting the kernel density.

For a third estimator, we use a parametrisation with seven parameters: (*π*_0_, *π*_1_, *π*_2_, *τ*_1_, *τ*_2_, *σ*_1_, *σ*_2_), which parametrise a four-part bivariate mixture-Gaussian distribution over the (+, +) quadrant with PDF:

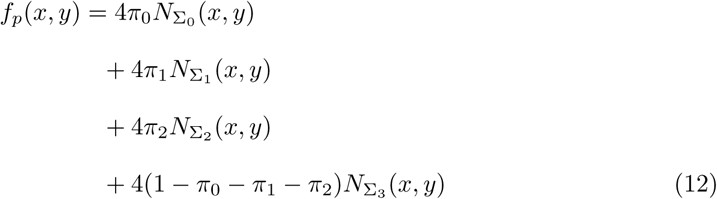

where *N*_Σ_(*x, y*) is the PDF of the bivariate normal distribution centred at the origin with variance Σ, the factor of 4 is due to to only unsigned *Z*-scores being used, and

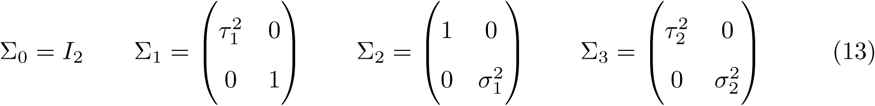

This model allows for a proportion *π*_1_ of study variables to be associated only with the trait of interest *P* (with *SD*(*Z_P_*) = *τ*_1_), a proportion *π*_2_ to be associated only with the second trait *Q* (with *SD*(*Z_Q_*) = *σ*_1_), and a proportion (1 − *π*_0_ − *π*_1_ − *π*_2_) to be associated with both (*Var*(*Z_P_, Z_Q_*) = Σ_3_). We allow different values of *σ*_1_, *σ*_2_ and *τ*_1_, *τ*_2_ to allow for potentially different reasons for shared (both *P* and *Q*) and independent (*P* XOR *Q*) associations. We make the not-generally-true assumption that *Z_P_* ⫫ *Z_Q_* in the final category in order to keep the model simple and avoid fitting covariances to absolute Z-scores. Maximum-likelihood estimates of parameters can be obtained using an E-M algorithm (Dempster *and others*, 1977). Given fitted parameters 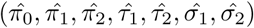, the denominator of (3) can be written:

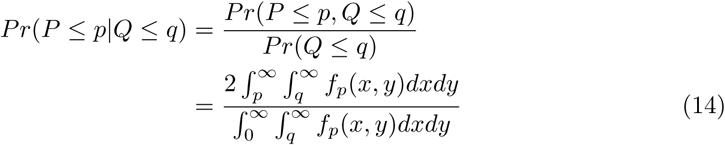

We denote the estimate of cFDR obtained using this method as

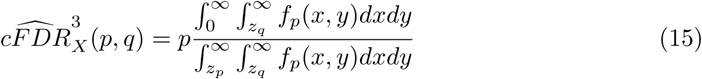

where *X* in this case is the set of points used in the estimation of parameters 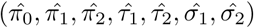.

Figure 2 shows the approximate behaviour of L-curves with each cFDR type. Notably, they are smoothest with 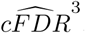, although they are constricted to a particular shape.

**Figure 2:**
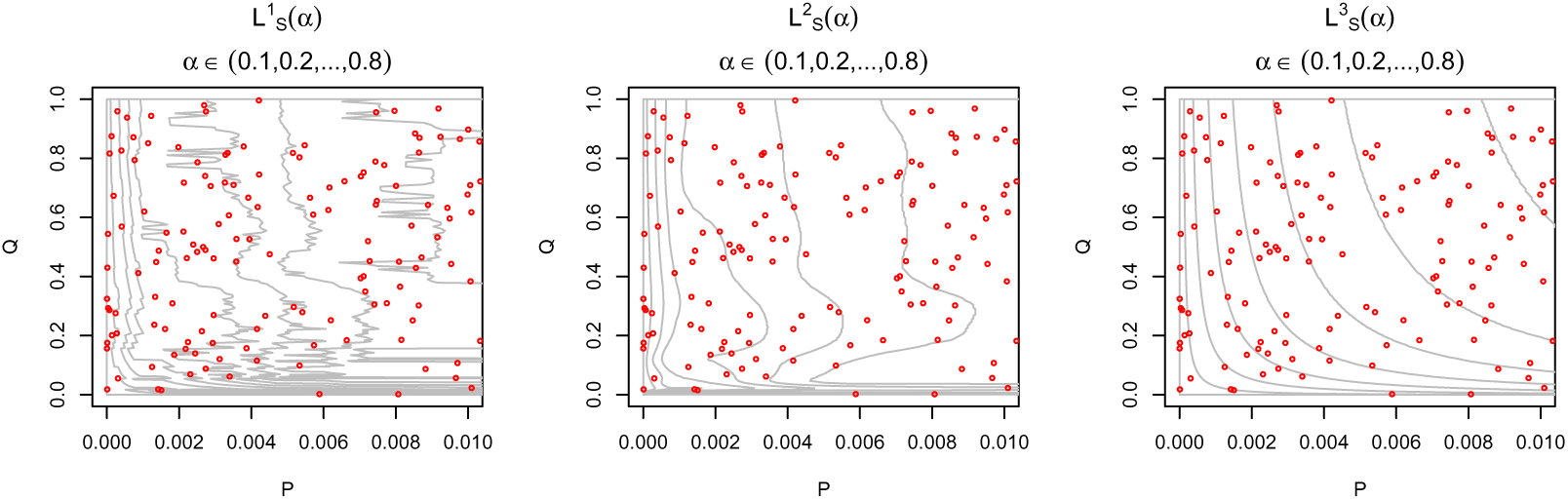
Forms of regions 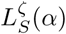 for *ζ* ∈ 1, 2, 3 (CDF, KDE, and modelled estimator respectively) for various values of *α*. Gray curves show rightmost border of regions, and red circles show datapoints in *S*. Curves (and hence rejection regions) are smoothest when *ζ* = 2, but are least responsive to local changes in data density. All regions tend to widen with lower *Q*.

Estimator 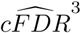 models the global distribution of *P, Q*, allowing simple approximation of the ‘local’ false-discovery rate:

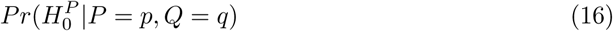

which can be more useful than the 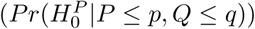. These quantities cannot be readily estimated by counting points in the manner of 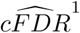.

### 3.2 Local cfdr

An important advantage of the parametrisation approach in 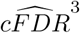 is the ability to estimate the quantity 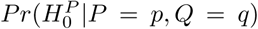 (which we will denoted 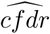 following the FDR/fdr convention (Efron *and others*, 2008)). We will denote by 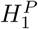 the complement of 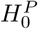 and *f*_1_(*p, q*) and *f*_0_(*p, q*) the density functions for 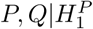 and 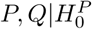 respectively.

We show here that the function *cfdr*(*p, q*), if known, can be used to generate optimal rejection regions for 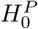, in the sense of regions *R* which maximise 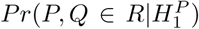 = 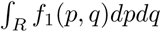 for a fixed value of 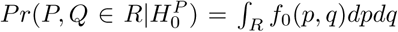 (where 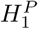 is the general alternative hypothesis for *P*). We briefly show that such regions *R* are regions inside contours of 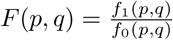.

Let *H* be the region of the unit square on one side of a contour *F* (*p, q*) = *c*, such that for *p, q* ∈ *H* ⇔ *F*(*p, q*) ≥ *c*, and suppose 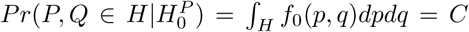 and that *f*_0_ > 0 on the interior of the unit square. Let *H′* be some other region for which 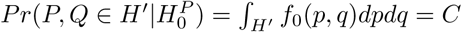. Now

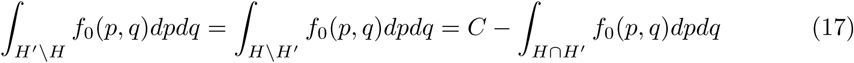

and since *F*(*p, q*) ≤ *c* for (*p, q*) ∈ (*H′* \ *H*) ⊆ *H′*, and *F*(*p, q*) ≥ *c* for (*p, q*) ∈ (*H* \ *H′*) ⊆ *H*, we have

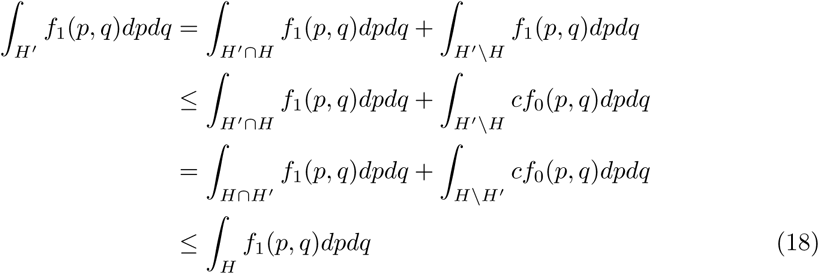

so amongst all regions *H′* with 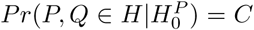 the region *H* maximises *Pr*(*P, Q* ∈ 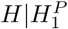). This means that if *H* is used as a rejection region, it will have the greatest power amongst all rejection regions corresponding to a type-1 error rate of *C*.

Since

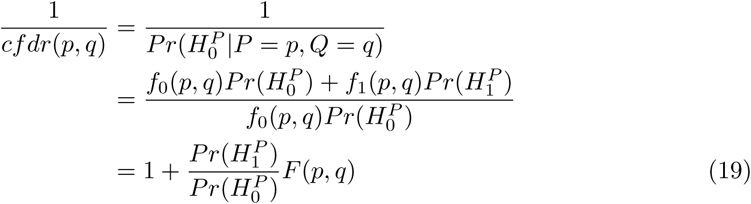

contours of *F* (*p, q*) are also contours of *cfdr*(*p, q*). Hence for any test statistic *k*(*p, q*) and corresponding threshold *α_k_* for which the rejection procedure *k*(*p, q*) ≤ *α_k_* controls the type 1 error at *α*, the test statistic *cfdr*(*p, q*) will be the most powerful.

Generally, as 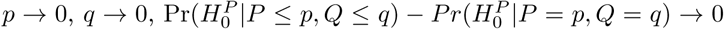, so the value *cF DR*(*p, q*) is a reasonably good approximation of *cfdr*(*p, q*), and contours of the two functions are similar. However, the argument above suggests that estimators for *cfdr*(*p, q*) may outperform estimators for *cFDR*(*p, q*) in hypothesis testing. In model 12, null variables (for which *Z_P_* ~ *N*(0, 1)) are in the first and third parts, meaning that under the model (writing *f*_0_ as a function of *z_p_, z_q_* for convenience)

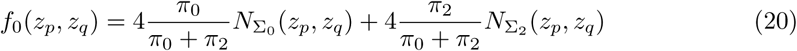

allowing the estimator

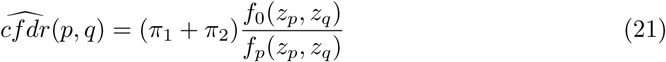

### 3.3 Estimation of 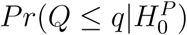

The FDR controlling method detailed in Section 2 does not require that 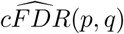 be a conservative estimator of 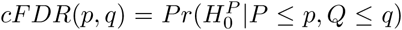. We can thus also estimate the quantity (5) as

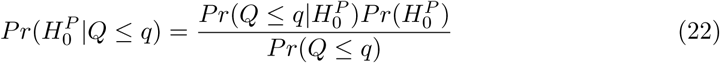

rather than approximating it upward to 1, as done previously. This is useful as the approximation upward to 1 is least accurate (and most conservative) when 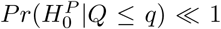, and it is in this situation that we can best use the event *Q* ≤ *q* to help reject 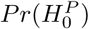. There is no need to estimate the quantity 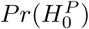 in the hypothesis-testing setting, as it is constant in all 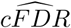 values and serves only as a scaling factor.

Any of the three estimators 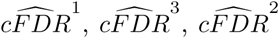 can be augmented by including an estimate of the quantity 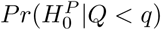, to improve accuracy. We use the estimate

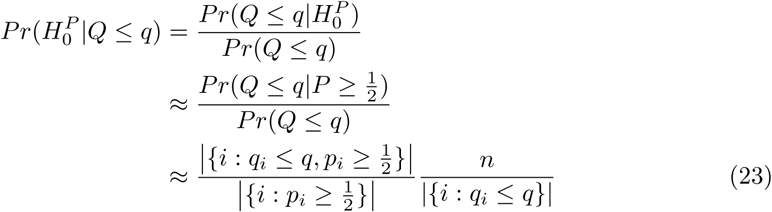

This can be used as a multiplicative factor in any of the estimators above. We denote

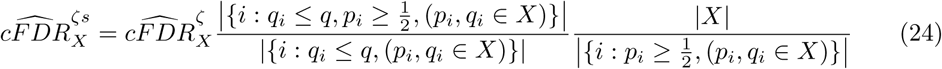

for *ζ* ∈ {1, 2, 3} as the ‘adjusted’ cFDR estimate after inclusion of this factor. Estimation of the density 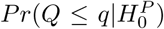 is also required for FDR control; namely in estimating the distribution of 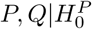 in order to integrate it over L. As described in our earlier paper Liley and Wallace (2018) we use a different strategy in this instance, by assuming a parametrisation of the distribution of 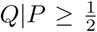 and estimating parameters using an expectation-maximisation algorithm. The parametrisation is far simpler to integrate over, so we prefer this method for integrating over regions *L*. If we used this parametrised estimate of 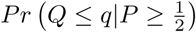 in place of the empirical CDF in formula (23), then since the CDF-based estimator of *Pr*(*Q* ≤ *q*) is at least 1/*n*, the estimate of 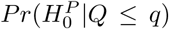 and hence the estimate of 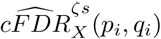 can be arbitrarily small for low *q_i_*, whatever the value of *p_i_*, especially if the true distribution of 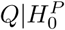 is not mixture-normal. We thus prefer the counting-points method for computing the quantity in the context of estimating cFDR values, and the parametrisation for integrating over regions *L*.

In both the multiplicative factor for 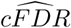 and integration over *L*, we approximate

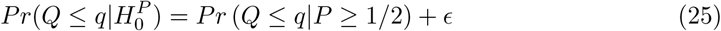

We note that (assuming 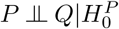)

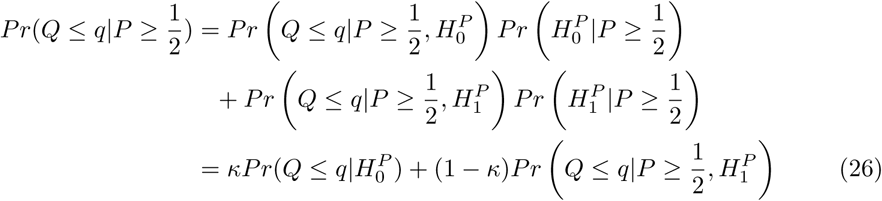

where 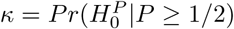, the approximation is consistent in that *κ* → 1 ⟹ ϵ → 0. A larger threshold on *P* than 1/2 can and should be used if sufficient points are available to estimate the density on the RHS of equation (25).

We would expect that *κ* →1 as sample sizes tend to ∞. For fixed finite sample sizes, the sign of *ϵ* is the sign of

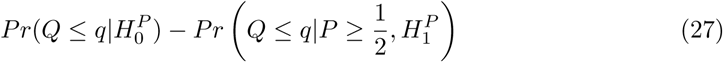

This quantity is difficult to make any judgements on; a tendency for common effects to both studies to be stronger than associations unique to the conditional trait would increase *ϵ*, but a tendency for effect sizes to be correlated at these common effects would decrease *ϵ*.

## 4 Assessment of performance through simulations

In order to assess the performance of each method above, we simulated a series of datasets and estimated the FDR of each method in each dataset. In each simulation, we generated a set of values *p_i_, q_i_, i* ∈ 1..*n* from random variables *P, Q*. We considered an extensive range of underlying parameters governing the distributions of *P_i_, Q_i_*. We used an identical simulation protocol to that used in (Liley and Wallace, 2018), reproduced in appendix 1.1. In general, each dataset contained 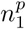 associations uniquely in the study 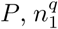 association uniquely in *Q*, and 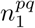 with both. We used various alternative distributions (normal, t (3df) and Cauchy) for simulating distributions of p-values under the alternative. We selected parameters from continuous distributions where possible to evaluate how power changed continuously.

For each simulation, we defined the ‘true detection rate’ (TDR) as the proportion of potential true discoveries made:

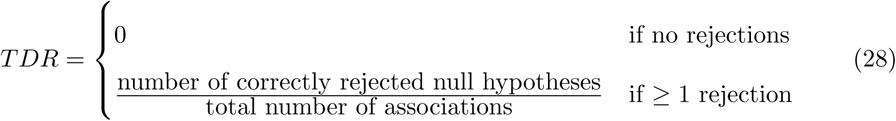

so the power to reject 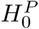 for a randomly-chosen true association is *E*(*TDR*). We controlled the FDR at *α* = 0.1.

We examined how power/TDR varied with the value of 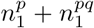, the total number of variables associated in the study of interest *P*. This displays the behaviour of estimators over the range of proportions of association variables from no associations at all 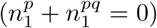 to a large number of associations.

In these simulations, we assume 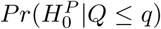 was known for all *q* for the purposes of integrating over *L* (that is, when we used the true distribution under which the data were simulated). An analysis of the effect of approximation of 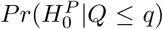 on FDR is made in Liley and Wallace (2018). All estimates of 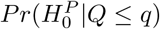 in the computation of 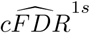, 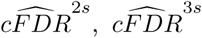 were made empirically throughout. We consider the performance of 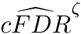 with *ζ* ∈ {1*s*, 2, 2*s*, 3, 3*s*} compared to *ζ* = 1

We compared numerical and graphical analyses of data using paired Wilcoxon rank-sum tests when comparing TDR values between different estimators. In comparing TDR between estimators, all Wilcoxon tests had 90% power to detect a difference between *Pr*(TDR with estimator A > TDR with estimator B) and *Pr*(TDR with estimator B > TDR with estimator A) of 5% at *p* < 0.05. P-values are not shown but ‘difference’ refers to *p* < 1 × 10−^3^ and ‘no difference’ refers to *p* > 0.1.

We compared the power of 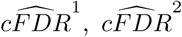, and 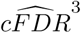 (figure 3) with each other and with the power of the Benjamini-Hochberg procedure applied to values *p_i_* at the same FDR (which we term BH). Estimator 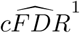 had the highest overall power, followed by estimator 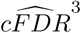 and method 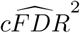. All method were more powerful than BH. When using a Gaussian alternative distribution, power was similar between estimators 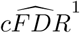 and 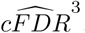, which was unsurprising given the bivariate Gaussian model used in 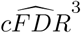. We found similar results when controlling at a more stringent FDR (*α* = 0.01), shown in supplementary figure 9

**Figure 3:**
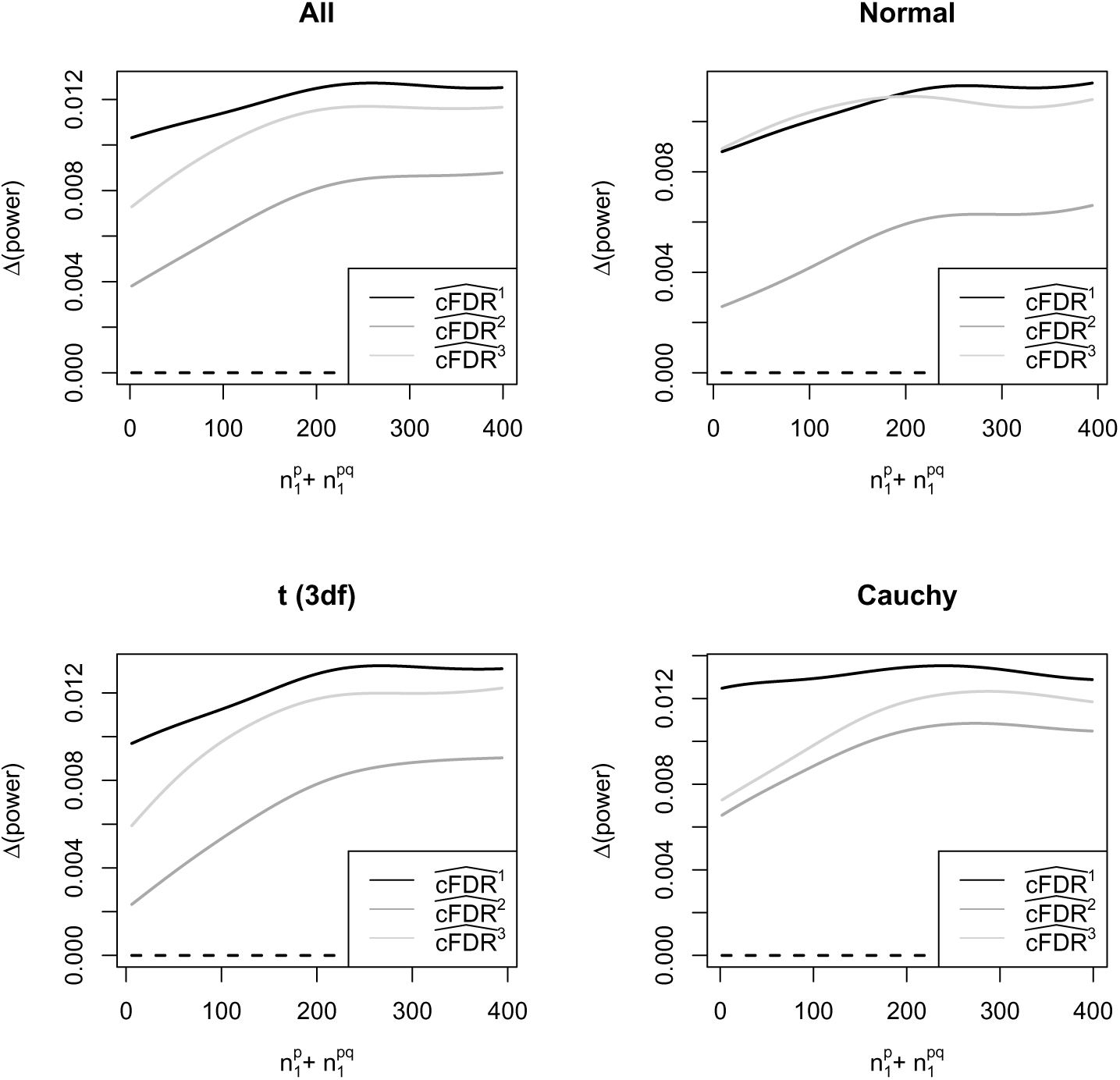
Relative power of each cFDR type (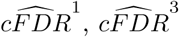, and 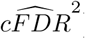) compared to BH, plotted against 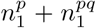 and subdivided by alternative distribution type (normal, t (3 df), Cauchy, and all combined with frequency 1/3). Parameters drawn from continuous distributions (see appendix 1.1. Gaussian smoothing is used with a kernel width of 1/8 of the x-axis range.

In all three estimators, the inclusion of the estimate of 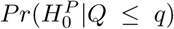 detailed in Section 3.3 improved the power of the procedure substantially (figure 4)

**Figure 4:**
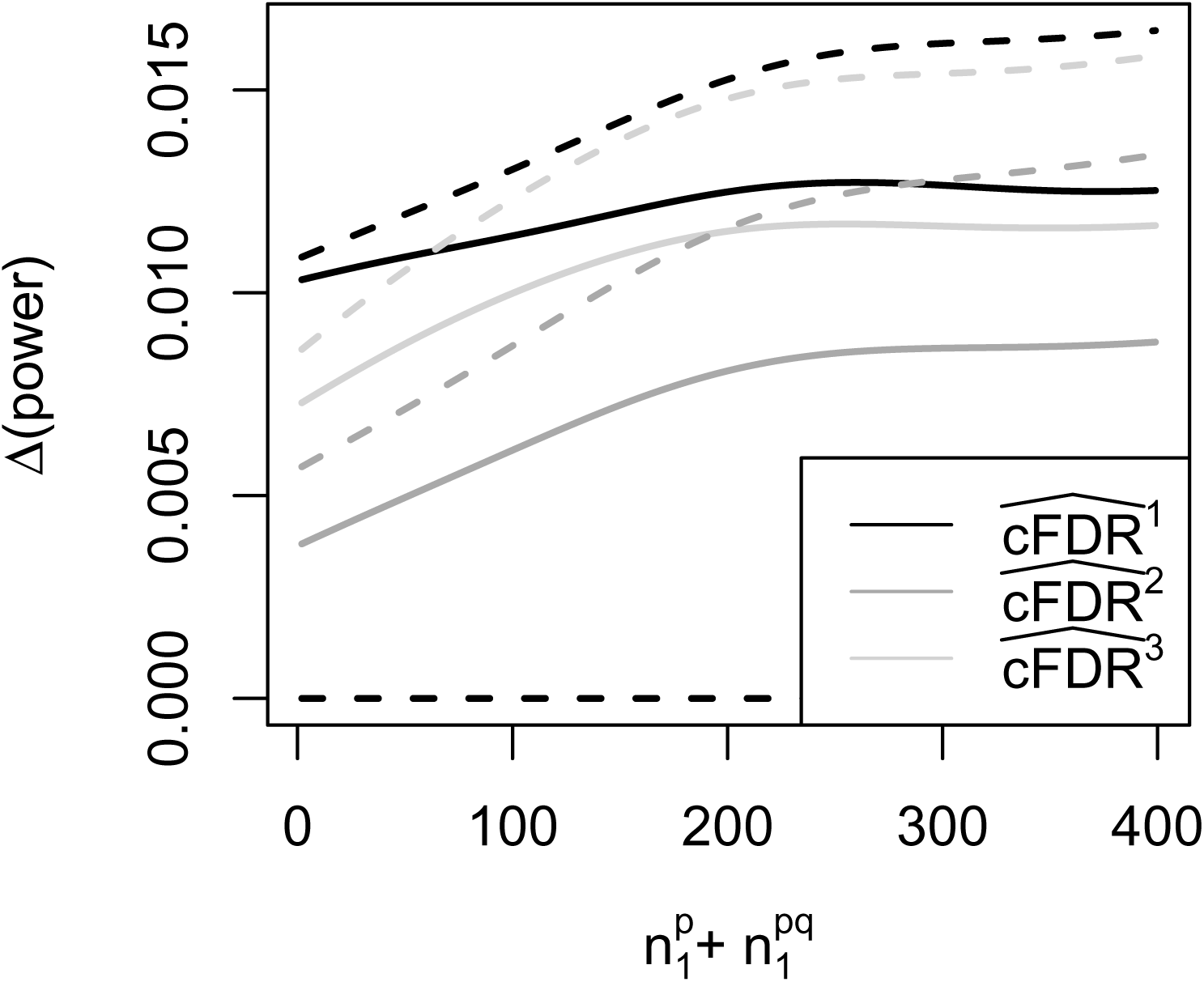
Difference between power of each cFDR type (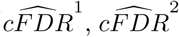, and 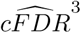) with and without inclusion of the estimate of 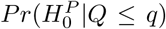 (that is, 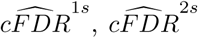, and 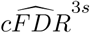), all compared to BH (the power of the Benjamini-Hochberg method applied to values *p_i_*. Dashed lines show ∆(power) for 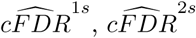, and 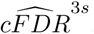.

Power was substantially higher using standard 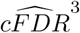 as opposed to 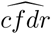, to the extent that that 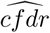 was less powerful for hypothesis-testing than the Benjamini-Hochberg method applied to p-values (figure 5). This was due to the chaoticity of contours derived from cfdr. An example of this is shown in figure 6. FDR control was maintained for all 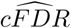 estimators (data available in github repository in ‘Data availability’).

**Figure 5:**
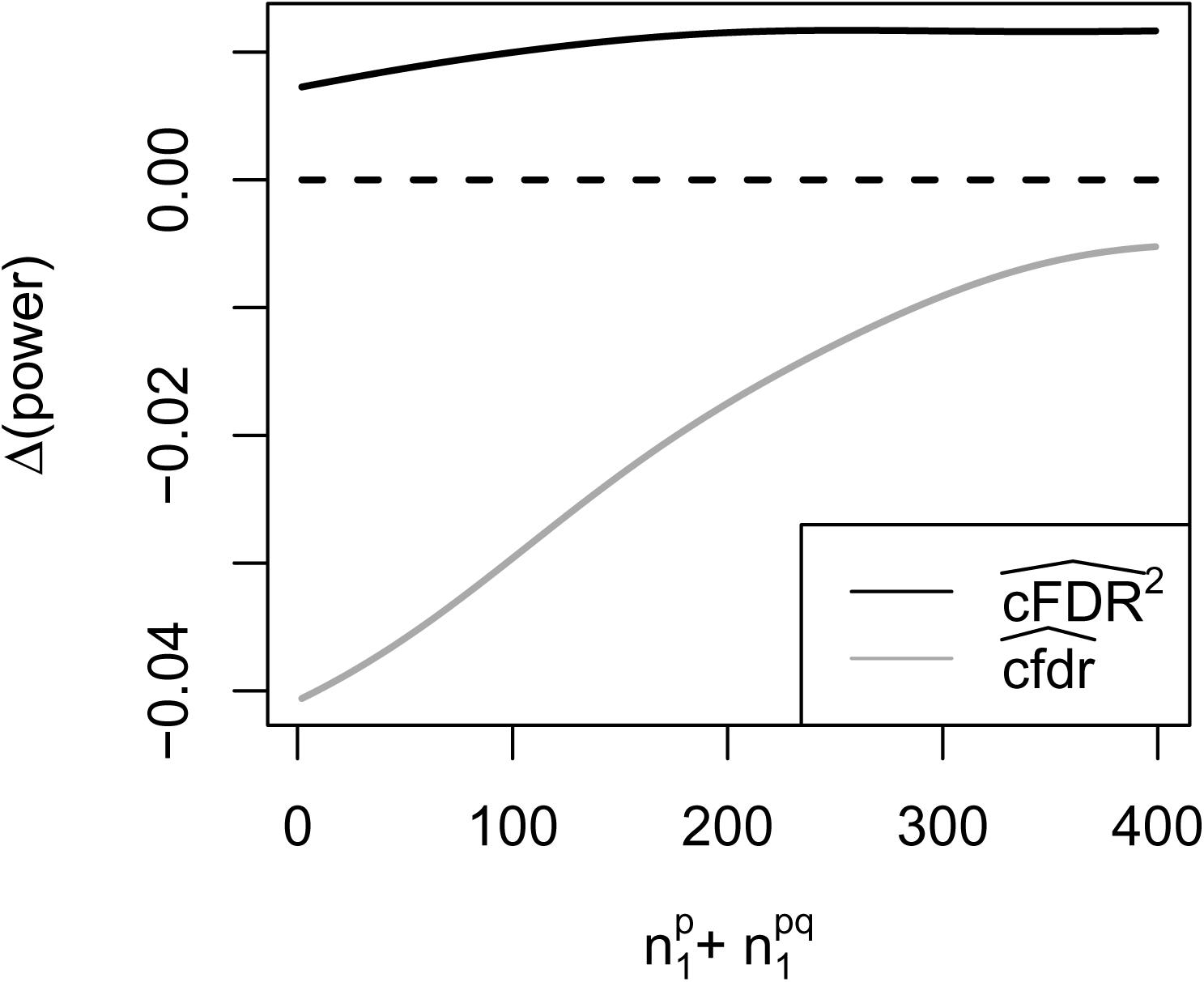
Power of local cfdr estimate 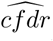 compared to cFDR estimate 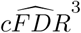, relative to the power of the Benjamini-Hochberg method applied to p-values. The local cfdr estimate performs poorly.

**Figure 6:**
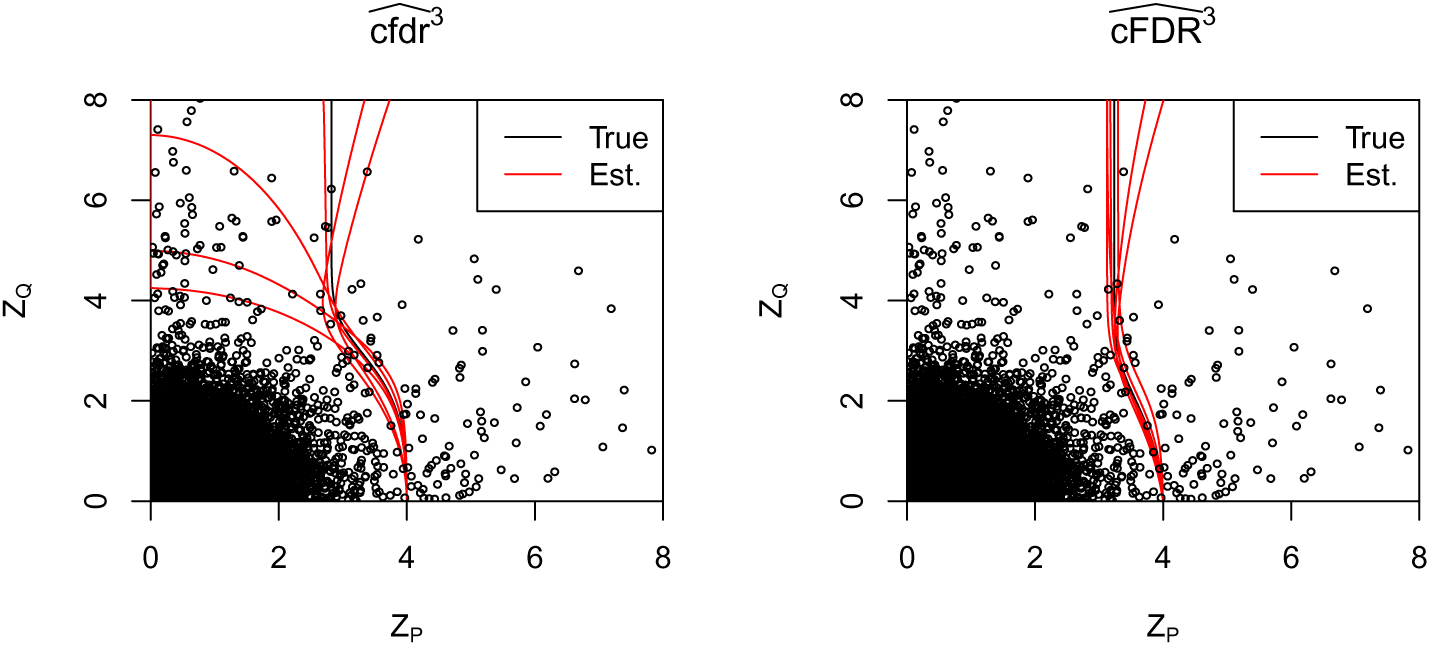
Contours of 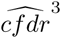 (‘local’ PDF-based estimator) and 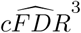 (standard CDF-based estimator) evaluated on the same data. Data were repeatedly simulated (six times) under the true parameter set (*π*_0_, *π*_1_, *π*_2_, *σ*_1_, *σ*_2_, *τ*_1_, *τ*_2_) = (0.94, 0.02, 0.02, 3, 3, 3, 3), with *n* = 10^5^ variables. For each simulation, MLE parameters were re-estimated. Red curves show contours of 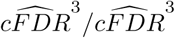 passing through (*Z_P_, Z_Q_*) = (4, 0), and black curves contours derived from the true (rather than estimated) parameters. Contours for 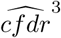 are far more chaotic, leading to loss of power despite theoretical optimality. Black points show a realisation of simulated *Z_P_, Z_Q_*.

## 5 Conclusions and recommendations

Based on our theoretical and simulated findings, we recommend the use of the 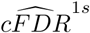 estimator for optimal power in hypothesis testing.

We recommend other methods in certain cases. If the alternative distribution of *Z* scores is likely to be normal (eg, *d* = 1 in table 1) or known, then cFDR estimator 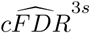 (or an equivalent with the known distribution type in place of the Gaussian functions in equation (12)) is of comparable power to 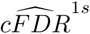 (see figure 3, panel ‘Normal’) and leads to smoother rejection regions and a less chaotic estimator. It also allows computation of 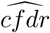 although we do not recommend its use for hypothesis testing unless there is great confidence in the approximation of the distribution of *P, Q*.

If a conservative estimate of the value 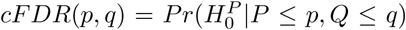 is needed, 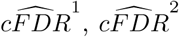, or 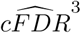 should be used in place of 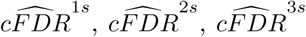. The inclusion of the estimate of 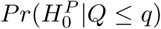, which is based on counting points, means that the continuity of 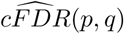 on (*p, q*) ∈ (0, 1)^2^ is sacrificed when using 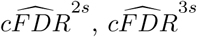 in place of 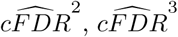. Hence if continuity of the estimator in *p, q* is required, 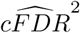 or 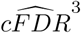 should be used rather than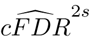 or 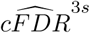.

## 6 Demonstration of method on TWAS data

In this section, we demonstrate the performance of the cFDR method on a real-world dataset from two TWAS. TWAS is a method to attempt discovery of gene-trait associations, as opposed to the variant-trait associations found by genome-wide association studies (GWAS). A TWAS firstly uses GWAS and expression-quantitative trait locus (eQTL) data to predict mRNA expression levels in a given tissue for each individual, then compares these predicted expression levels across a trait of interest Gusev *and others* (2016).

We considered TWAS datasets for breast cancer (BRCA, Michailidou *and others* (2017)) and ovarian cancer (OCA, Phelan *and others* (2017)), containing tests for varying numbers of genes across 54 tissues. BRCA and OCA have considerable phenotypic overlap Greene *and others* (1984), and we may hope that summary statistics for one disease may be useful for leverage in association analyses of the other. We considered RNA-tissue pairs available in both datasets, restricting our analysis only to pairs in which RNA expression was predicted using data from the GTEx consortium, comprising a total of *n* = 80222 hypotheses.

Given the GWAS-scale dimensionality of testing, we chose a conservative FDR control level *α* = 1 × 10^−6^. We used 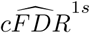 for cFDR estimation (and 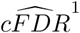 for contrast). We assigned folds according to genes, so expression levels for each gene were assigned a separate fold (for 11327 folds in total). Figure 7 shows z-scores for BRCA and OCA, and rejection regions for cFDR and p-value at the same level of FDR control *α*. The analysis of BRCA conditioning on OCA enabled 38 more gene-tissue association discoveries than when the p-value alone was used and notably nine more discoveries than when 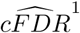 was used (724 for 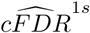 vs 715 for 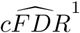 vs 678 for p-value). In the corresponding analysis for OCA conditioned on BRCA, both 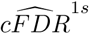 and 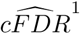 enabled four more discoveries than when using p-values alone (310 vs 306). Subjectively, rejection regions from 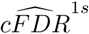 appear more natural than those from 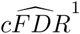 or from p-values alone, adapting to the joint distribution of Z-scores without requiring any prior hypothesis of joint association.

**Figure 7:**
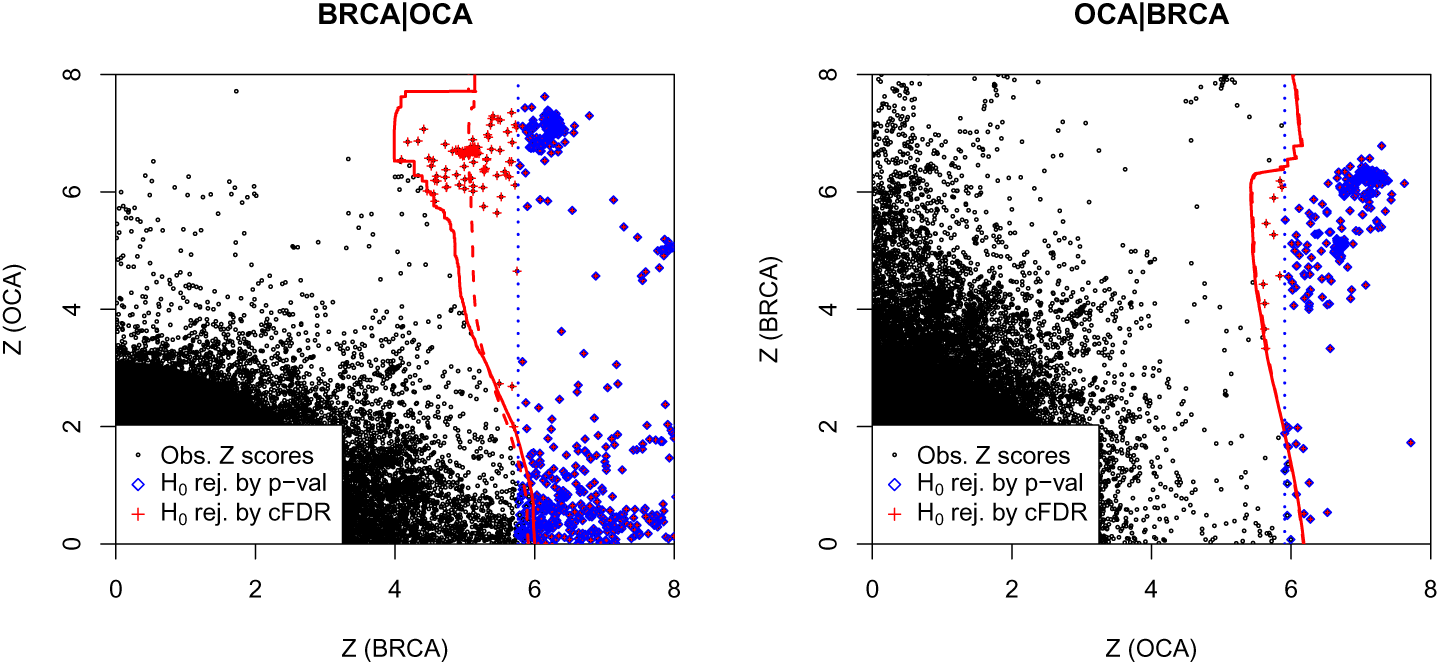
Z scores from TWAS on breast cancer (BRCA) and ovarian cancer (OCA) for a range of genes in 53 tissue types. Red points (crosses) show rejections using 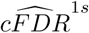, blue points (diamonds) using p-value alone, controlling overall FDR at *α* = 1 × 10−^6^. Red (solid) and blue (dashed) lines indicate approximate rejection regions for 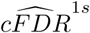 and p-values respectively. Dashed red lines (underneath solid lines on right panel) denote approximate rejection regions for 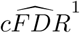

We found that the relative number of discoveries with the two methods varied considerably with *α* (figure 8). This observation is specific to this application, and should not be generalised to other TWAS nor to any assumptions on the behaviour of the cFDR. We also do not endorse the universal use of any FDR threshold (including the one used above) in TWAS, and especially note that *α* should not be chosen to maximise the number of discoveries after an analysis such as that in figure 8. A study-appropriate threshold should be determined *a priori* by consideration of the relative cost of type 1 and 2 errors.

**Figure 8:**
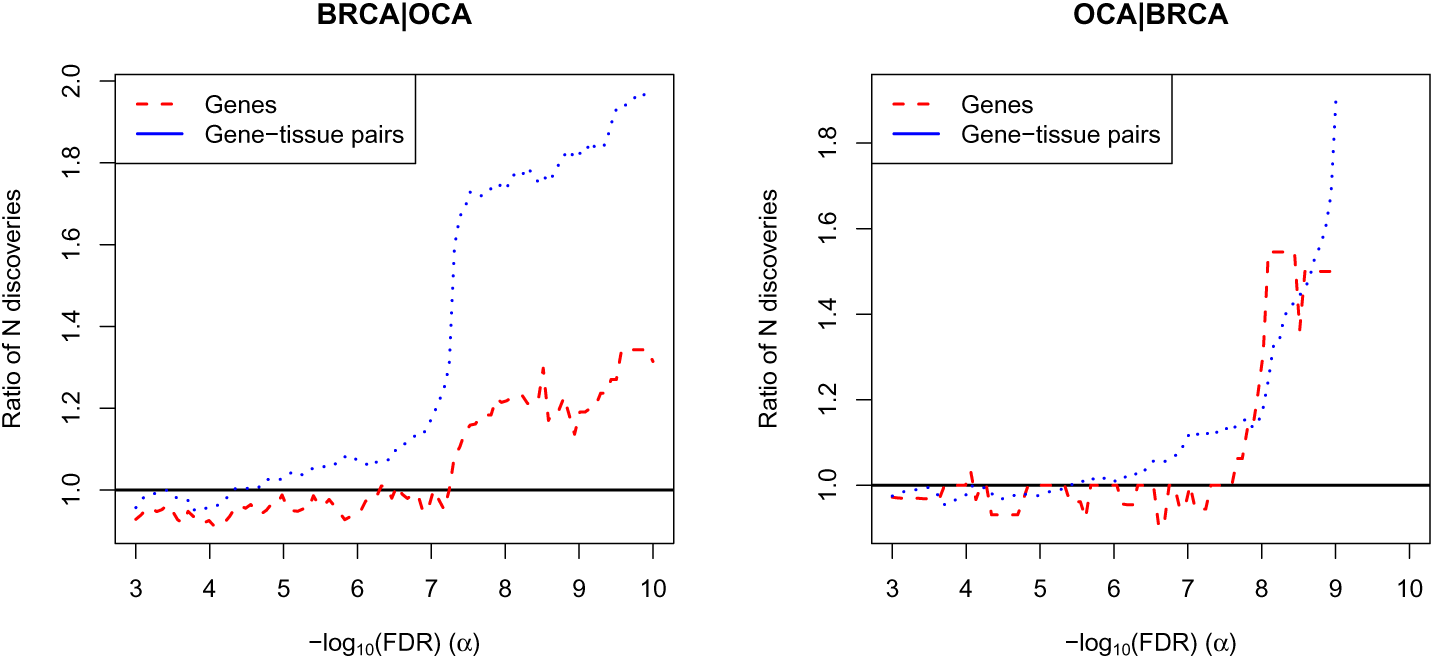
Ratio of new discoveries using 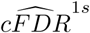 and new discoveries using p-value at varying *α*. New gene-disease associations are shown in red, new gene-tissue-disease associations in blue. Both trend upwards with decreasing *α*.

This section is intended to be demonstrative rather than exploratory and we do not claim that putative cFDR-based discoveries are novel. Since this demonstration only involves a single dataset, any judgement about the appropriateness of cFDR to a particular application should be made using the results of the simulations above, rather than the performance on this dataset.

## 7 Discussion

There are two main ideas in this work; firstly, the use of parametric or KDE-based estimators to address the discontinuity exhibited by the empirical estimator 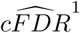 and secondly the augmentation of any estimator by inclusion of an estimate of 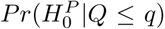. While they were generally less powerful than 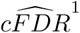, parametric and KDE-based cFDR both reduced the chaoticity of the estimator around extremal points. An important pragmatic finding of this work was that the standard non-parametric cFDR is hard to improve on. We showed this both in a theoretical sense, in that it resembles the optimal cfdr, and in an empirical sense in the observation that neither of the alternative estimators were stronger.

The parametric cFDR estimator 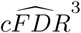 was slightly more powerful than the non-parametric estimator 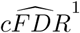 when the underlying distributional assumption (Normal alternative) was correct. The difference in power in this case was small, and 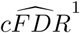 was considerably more powerful when the distributional assumption was false. As *n* → ∞, we expect that 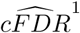 will be more powerful than 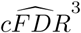 if distributional assumptions are false, and both will have equivalent power if they are true (since L-regions associated with 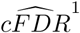 will converge towards the ‘true’ L-regions associated with the underlying *P, Q* distribution, and L-regions associated with 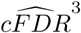 will converge towards those associated with the best normal approximation to it).

The parametric cFDR estimator has several theoretical advantages. First, it invites an extension to higher-dimensional spaces, conditioning on two traits rather than one, which is an interesting avenue for future work. Second, it allows computation of the ‘local’ cfdr, which defines a different ordering on hypotheses than the cFDR. This contrasts with the one-dimensional case, where fdr estimates, FDR estimates, and p-values are generally monotonic (Efron *and others*, 2008). Although the cfdr corresponds to a theoretically optimal decision boundary for rejecting 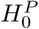 if estimated accurately, we found that the estimate was too chaotic to be of general use in hypothesis testing with *n* < 10^4^.

It is possible that other estimators of cfdr may be more stable. However, we suspect that similar problems are likely to be encountered, as PDFs are generally harder to estimate than CDFs, even in the univariate setting (Efron *and others*, 2008); heuristically, to estimate the *PDF* of *P, Q* at a point (*p, q*), either only the small proportion of the data local to (*p, q*) can be used, or strong parametric assumptions must be made to allow distant points to affect the estimate. We note that cfdr performed poorly even when distributional assumptions were correct and the underlying distribution was well-behaved (Normal) and had only seven degrees of freedom. These artificial conditions are far simpler than those encountered in ‘real-world’ problems, and we conclude that while the cfdr may remain a useful Bayesian quantity in assessing the posterior probability of association for an interesting variable, no estimator of cfdr is likely to be of use in hypothesis-testing at these scales (*n* < 10^4^). Other estimators (for instance, that of Zablocki *and others* (2014)) may be effective at higher *n*.

Inclusion of an estimate of the quantity 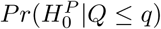 in the cFDR estimate improves the power over existing methods, and the accuracy of the estimate, at the cost of asymptotic conservatism. The assumption 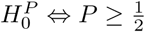 is not ideal, since the sign of ϵ in equation (25) cannot usually be confidently determined, and the estimation is not consistent as *n* → ∞ (although it is if sample sizes → ∞). Estimates of 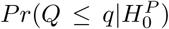 may thus be biased. Alternative methods for estimating 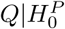 may include examining the distribution of *Q* amongst P-values reaching a higher threshold on *P* (for instance, assuming 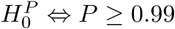) or amongst a set of ‘control’ variables known to be in 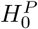 and typical of all variables in 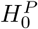. However, in Liley and Wallace (2018), we found that the estimate tended to be adequate in the context of integrating over regions *L*.

In summary, our methods add to the body of methods for co-analysis of omics data. Estimating 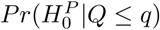 improves the power of cFDR estimators, and parametric estimators enable greater flexibility in their use. We present empirical arguments that although the cFDR is not a theoretically optimal discriminator, it may be pragmatically the best available. These results support confidence in the robustness of the 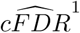 estimator generally and associated FDR-controlling procedures, even at relatively small values of *n*.

## 8 Software

Software is available as an R package at https://github.com/jamesliley/cfdr. All data and a pipeline to deterministically generate all plots and results in this paper can be found at https://github.com/jamesliley/cfdr_estimation_pipeline

## 9 Supplementary Material

1. Description of our protocol for simulations
2. Supplementary figure: comparison of cFDR estimators at *α* = 0.01

**Figure 9:**
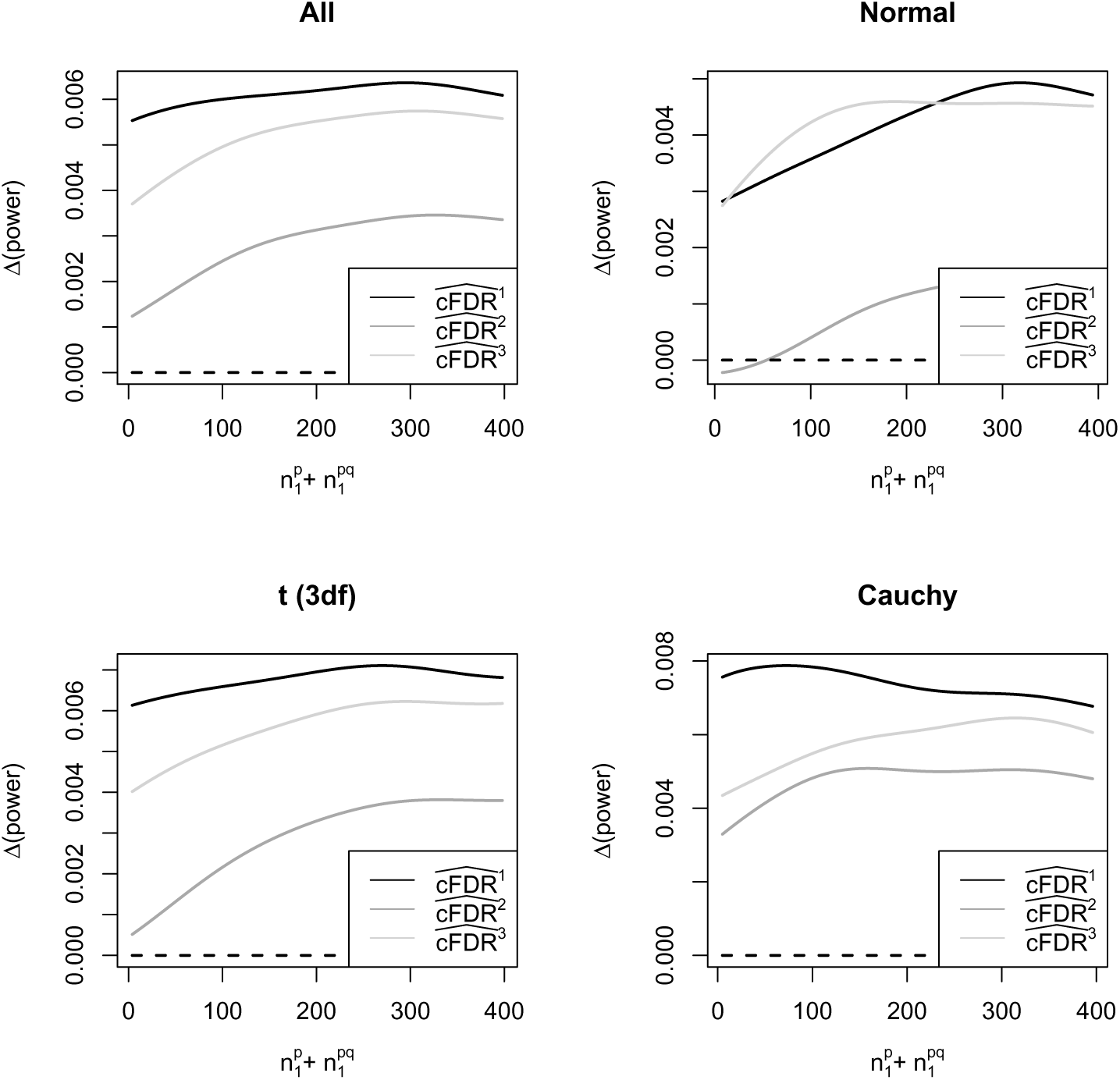
Analogous to figure 3 with *α* = 0.01 instead of *α* = 0.1. Relative power of each cFDR type (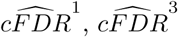, and 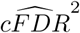) compared to power of the Benjamini-Hochberg method, using FDR control method 3b, plotted against 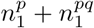 and subdivided by alternative distribution type (normal, t (3 df), Cauchy, and all combined with frequency 1/3). Parameters drawn from continuous distributions (see appendix 1.1). Gaussian smoothing is used with a kernel width of 60.

## Acknowledgments

CW is funded by the Wellcome Trust (WT107881) and the MRC (MC UU 00002/4). JL is funded under by WT107881 and for part of this work was on the Wellcome Trust PhD programme in Mathematical Genomics and Medicine, funded by the NIHR Cambridge BRC. The funders had no role in study design, data collection and analysis, decision to publish, or preparation of the manuscript.

## General notes for notation

1. *P, p* refer to study under investigation, *Q, q* to study for conditioning on. *P* and *Q* are also used to refer to these two studies as opposed to the random variables corresponding to p-values.
2. *i* and *j* index hypotheses and variables corresponding to hypotheses, (*i*) generally meaning the *i*th smallest
3. *S* refers to the set (*p*_1_, *q*_1_), (*p*_2_, *q*_2_), …, (*p_n_, q_n_*)
4. *k* indexes folds/subdivisions
5. *n* is the total number of hypotheses
6. *N* is the number of folds for cross-validation/resampling
7. *P, Q* are random variables, *p, q* are observations
8. *H^P^* is an indicator variable for association in study *P*. It can be considered in a Bayesian sense as a Bernoulli-distributed random variable, or a frequentist sense as a null hypothesis eg *H^P^* = 0.
9. v-values refer to values *v*(*L*) used for testing in some sense. The exact definition depends on the method used for FDR control.

## 1 Appendix

### 1.1 Simulation protocol

The distributions and variables we considered are shown in table 1. In general, our simulations took the form of: a fixed number 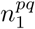 of variables associated in both *P* and *Q*, a fixed number of variables 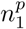 associated only with *p*, a fixed number of variables 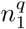 associated only with *Q* and the remainder of variables associated with neither *P* nor *Q*. We will refer to these four classes of variables as *C*_1_, *C*_2_, *C*_3_, *C*_4_. We designated that within each class of variables, *P_i_* and *Q_i_* were iid.

**Table 1:** Variables used in simulations

| Variable | Description | Point values | Sampling distribution for other values |
| --- | --- | --- | --- |
| $n$ | Total number of variables | $10^3, 10^4$ | $10^{U(3,4)}$ (rounded) |
| $n_1^{pq}$ | Number of variables assoc. with $P, Q$ | 0, 10, 200 | $U(0, 200)$ |
| $n_1^p$ | Number of variables associated with $P$ | 0,10,200 | $U(0, 200)$ (rounded) |
| $n_1^q$ | Number of variables associated with $Q$ | 0,10,200 | $U(0, 200)$ (rounded) |
| $s_p$ | Scale for distribution of $P_i$ (see below) | $\frac{3}{2}, 3$ | $U(\frac{3}{2}, 3)$ |
| $s_q$ | Scale for distribution of $P_j$ | $\frac{3}{2}, 3$ | $U(\frac{3}{2}, 3)$ |
| $d$ | Form of distributions | Normal, t (3df),<br>Cauchy (eq. prob.) | Normal, t (3df),<br>Cauchy (eq. prob.) |

We considered two types of simulations: either with all numerical parameters drawn uniformly from a set of fixed ‘interesting’ values, or with all numerical parameters drawn from continuous distributions. Our intent was to evaluate FDR values accurately at these ‘interesting’ points in the parameter space, and more roughly consider values in between these points. One such ‘interesting’ area of the parameter space was the scenario in which no variables were associated in study *P* (that is, 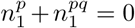). In some plots (namely when looking at how power varies with 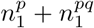) only simulations with parameters drawn from continuous distributions are used.

For variables in *C*_1_, *C*_2_, we set the distribution of *P_i_* (determined by *d, s_p_*) by first simulating Z scores:

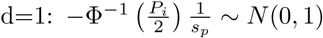

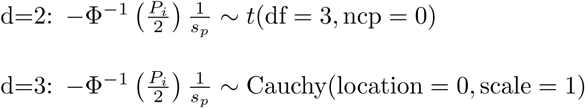

where 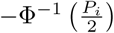 can be considered a *Z*-score corresponding to *P_i_*, and *s_p_* a scaling factor for the distribution. We set the distribution of *Q_i_* in *C*_1_, *C*_3_ similarly, with *s_q_* in place of *s_p_*. The values *p_i_, q_i_* for *i* ∈ *C*_4_ were sampled from *U*(0, 1).

Within each class *C*_1_ − *C*_4_ we simulated independent *P, Q*, although *P, Q* are still dependent when not conditioning on class. We also considered *s_p_* to be constant across *C*_1_ and *C*_2_, *s_q_* to be constant across *C*_1_ and *C*_3_, and *d* to be constant in each simulation.

We analysed each simulated dataset in parallel using each of the three estimators of 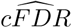. We also performed a standard Benjamini-Hochberg procedure on the values of *P* (without considering the values of *Q*) as a control. We further considered each estimator of 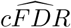 both with and without the adjustment due to estimation of 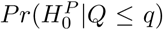. To estimate the null distribution of 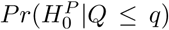 in order to integrate to generate values *v*(*L*), we considered both the true distribution and a mixture-normal distribution estimated from the distribution of 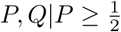 as per Section 3.3.

